# Differential remodeling of the electron transport chain is required to support TLR3 and TLR4 signaling and cytokine production in macrophages

**DOI:** 10.1101/742098

**Authors:** Duale Ahmed, David Roy, Allison Jaworski, Alex Edwards, Alfonso Abizaid, Ashok Kumar, Ashkan Golshani, Edana Cassol

## Abstract

Increasing evidence suggests that mitochondria play a critical role in driving innate immune responses against bacteria and viruses. However, it is unclear if differential reprogramming of mitochondrial function contributes to the fine tuning of pathogen specific immune responses. Here, we found that TLR3 and TLR4 engagement on murine bone marrow derived macrophages was associated with differential remodeling of electron transport chain complex expression. This remodeling was associated with differential accumulation of mitochondrial and cytosolic ROS, which were required to support ligand specific inflammatory and antiviral cytokine production. We also found that the magnitude of TLR3, but not TLR4, responses were modulated by glucose availability. Under conditions of low glucose conditions, TLR3 engagement was associated with increased ETC complex III expression, increased mitochondrial and cytosolic ROS and increased inflammatory and antiviral cytokine production. This amplification was selectively reversed by targeting superoxide production from the outer Q-binding site of the ETC complex III. These results suggest that ligand specific modulation of the ETC may act as a rheostat that fine-tunes innate immune responses *via* mitochondrial ROS production. Modulation of these processes may represent a novel mechanism to modulate the nature as well as the magnitude of antiviral versus inflammatory immune responses.

## Introduction

The innate immune system, including tissue macrophages, represent the first line of defence against invading microbial pathogens. Early recognition depends on a variety of pattern recognition receptors (PRRs), which detect evolutionarily conserved structures termed pathogen-associated molecular patterns (PAMPs)^1, 2^. Key players in this process are Toll-like receptors (TLR), which are capable of detecting a range of PAMPs from viruses and bacteria^1–5^. Among the best characterized are TLR3 and TLR4. TLR3 recognizes double stranded RNA (dsRNA), a common PAMP associated with viral infections^6^. TLR4 primarily recognizes lipopolysaccha r ide (LPS), the core component of the outer membrane of Gram-negative bacteria^7^. Both TLR3 and TLR4 differentially and dynamically modulate nuclear factor-κB (NF-κB) and interfero n regulatory factory 3 (IRF3) signaling following receptor engagement. While TLR3 activates NF-κB and IRF3 signaling *via* TIR-domain-containing adapter-inducing interferon-β protein (TRIF), TLR4 signals through both Myeloid differentiation primary response 88 (MyD88) and TRIF^1^. Differential activation of these signaling pathways plays a critical role in fine-tuning pathogen specific antiviral and antibacterial responses^1^.

Cellular metabolism has emerged as a key regulator of macrophage function. Metabolic reprogramming is required to meet the bioenergetic and biosynthetic demands of the cell and to drive effector functions^8–10^. Alterations in metabolites and other bioactive metabolic products have also been shown to activate and regulate gene expression, signal transduction and epigenetic profiles^9, 11–14^. Among the best characterized examples of metabolic reprogramming occurs following LPS stimulation^9, 14–18^. Almost immediately after TLR4 engagement, macrophages downregulate oxidative phosphorylation (OXPHOS) activity and dramatically increase glycolysis to support rapid ATP production^14, 15, 17–19^. This repurposing of mitochondrial function also increases reactive oxygen species (ROS) levels, which drives inflammatory cytokine production^9, 15, 17^. ROS production is driven by the combined effects of increased mitochondr ia l membrane potential and the oxidation of succinate by complex II of the electron transport chain (ETC)^15^ suggesting flux through the ETC may play a central role in this process. However, it is still unclear if the dynamic modulation of ETC complexes and increased ROS production contributes to signaling events following engagement of other TLRs and if different ia l reprogramming of these processes contributes to ligand specific immune responses.

Mitochondria are known to play an important role in innate immune responses against RNA viruses^20–23^. Recognition of cytosolic viral RNA by retinoic acid-inducible gene I (RIG-I)-like receptors (RLR) and their downstream processes have been shown to require the participation of mitochondrial antiviral signaling (MAVS), a mitochondrial outer membrane adaptor protein ^20, 24, 25^. MAVS acts as a scaffold and recruits effector molecules to the mitochondrial outer membrane leading to the activation of NF-κB and IRF3^20^. More recent studies have shown that mitochond r ia also contribute to antiviral signaling *via* MAVS independent mechanisms. Tal et al. found that mitochondrial ROS (mtROS) potentiates RLR signaling. This signaling is regulated by autophagy and clearance of dysfunctional mitochondria^26^. Alternatively, Yoshizumi et al. found that RLR-mediated antiviral responses are dependent on OXPHOS activity. This dependence is regulated by the mitochondrial fusion protein optic atrophy 1 (OPA1)^23^. These findings suggest that other aspects of mitochondrial function, independent of MAVS mediated scaffolding, may play a central role in facilitating antiviral responses.

While both TLR3 and RLR recognize and respond to virally derived RNAs, they signal through distinct transduction pathways to trigger antiviral immune responses^27, 28^. To date, the role of the mitochondria in driving TLR3-mediated responses in macrophages remains poorly understood. Previous studies have shown that engagement of TLR3 on hepatocytes and dendritic cells (DCs) causes a shift from OXPHOS towards aerobic glycolysis for energy production^29–32^. In DCs, this shift is driven predominately by the *de novo* production of type I interferons (IFN) and therefore is required to rapidly meet the increased energy demands of these activated cells^30–32^. Macrophages stimulated with high concentrations (10μg/mL) of the synthetic TLR3 agonist polyinosinic-polycytidylic acid (Poly(I:C) or PIC) have been shown to downregulate Complex I-associated ATP production under standard culture conditions^33^. However, the functional consequences of this ETC reprogramming has yet to be elucidated.

In the current study, we used murine bone marrow-derived macrophages (BMM) to evaluate how reprogramming of mitochondrial function contributes to TLR3 and TLR4 signaling and cytokine production and how glucose availability affected these responses. We found that modulation of flux through the ETC and associated ROS production plays a critical role in cytokine production following TLR engagement. This reprogramming is ligand specific and may have different ia l effects on the expression of individual cytokines (e.g. accumulation of mitochondrial vs. cytosolic ROS). Further, low glucose conditions resulted in differential reprogramming of mitochondr ia l function following TLR3 engagement. This reprogramming upregulated complex III expression and associated mitochondrial ROS production, which amplified inflammatory and antiviral signaling and cytokine production. Collectively, these findings suggest that the ETC may act as a selective rheostat of innate immune responses that differentially regulates ligand specific responses based on nutrient availability.

## Results

### Differential production of pro-inflammatory and antiviral cytokines in PIC-and LPS-stimulated BMM

Despite activating the same transcription factors (e.g. NF-κB and IRF3), signaling through TLR3 and TLR4 are associated with distinct inflammatory and antiviral cytokine profiles. To evaluate these differences in our model system, we stimulated BMM with PIC (10ng/ml and 10μg/ml) or LPS (100ng/ml) for 18 hours and assessed inflammatory (TNF-α, IL-1β, and IL-6) and antiviral (IFN-α, IFN-β and CXCL10) cytokine production in culture supernatants. PIC concentrations were selected to emulate responses in early (low levels of virus) and late stages of infection (high levels of virus), where Lin et al. found that only high (≥ 10μg/ml) concentrations of PIC can induce robust inflammatory cytokine production^34^. The LPS concentration was selected based on its ability to repurpose mitochondrial function to support ROS production^15^. As previously reported^15^, LPS induced a strong inflammatory response, produced intermediate levels of IFN-β and CXCL10 and no IFN-α (Figure 1). Alternatively, stimula t io n with low concentrations of PIC induced low levels of antiviral and inflammatory production. Increasing the PIC concentration (10μg/ml) significantly increased both inflammatory and antiviral cytokine production (Figure 1).

**Figure 1:**
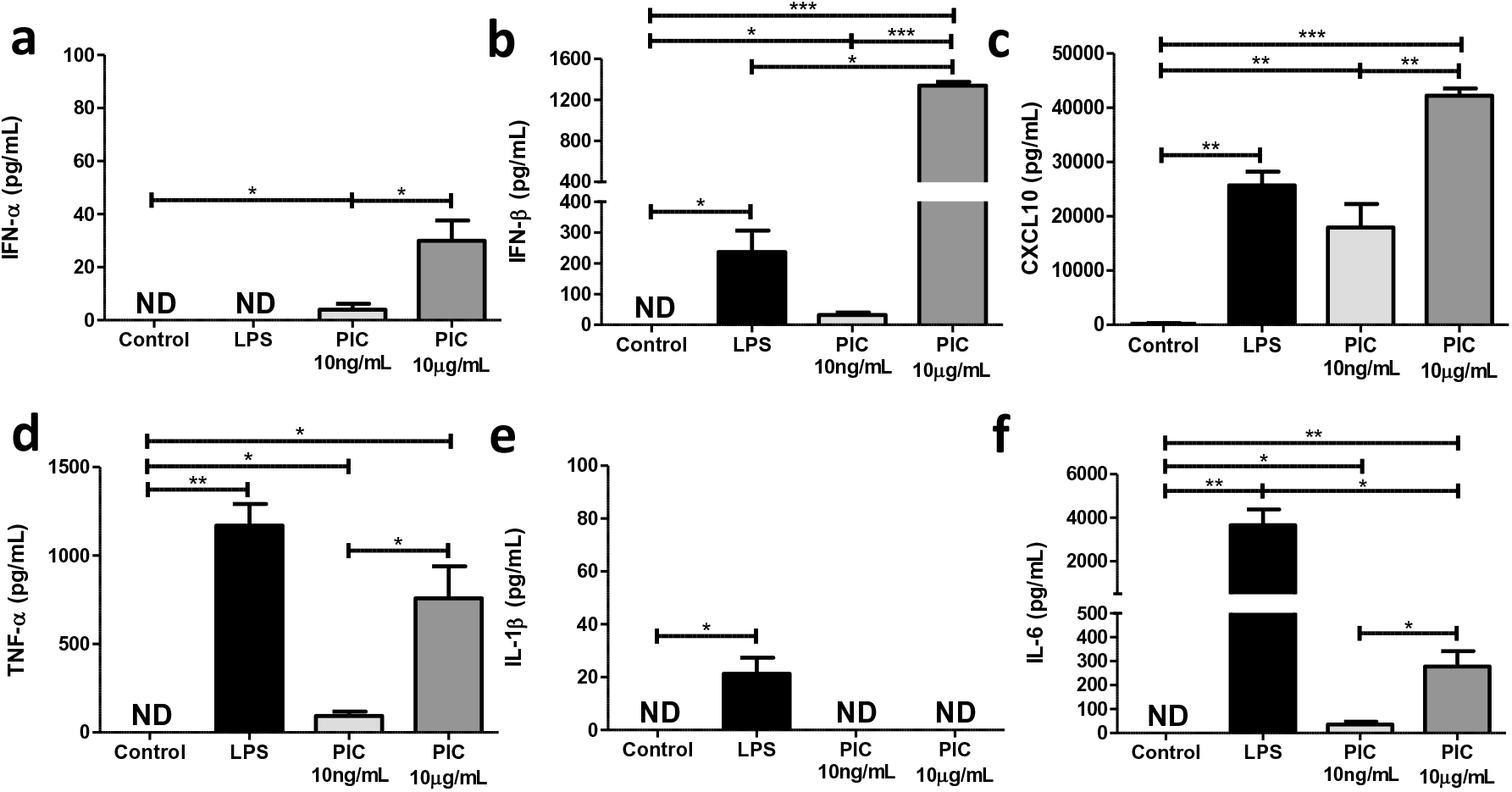
High, but not low, concentrations of Poly(I:C) are associated with pro-inflammatory cytokine production. Bone marrow-derived macrophages (BMDMs) were treated with either 100ng/mL lipopolysaccharide (LPS), 10ng/mL or 10μg/mL Poly(I:C) (PIC) for 18 hours. Supernatant was collected and assessed for antiviral (IFN-α, IFN-β, CXCL10) **(a-c)** and pro-inflammatory (TNF-α, IL-1β, IL-6) **(d-f)** cytokine expression. Data represents mean ± SEM of four individual mice (*p < 0.05, **p < 0.01, and ***p < 0.001).

### BMM stimulated with low versus high concentrations of PIC differ in their ability to ramp up glycolytic activity under stress

Next, we evaluated the differential effects of LPS and PIC stimulation on cellular metabolism using the Seahorse extracellular efflux analyser. As described above, BMM were stimulated with LPS (100ng/ml), low (10ng/ml) or high (10μg/ml) concentrations of PIC for 18 hours. Changes in proton efflux rate (PER) at baseline and in response to rotenone + antimycin (ROT/AA) and 2-deoxyglucose (2-DG) injections were used to evaluate changes in glycolytic parameters. As previously described^9, 35^, LPS stimulation increased PER levels at baseline, increased the %PER derived from glycolysis and decreased the mitoOCR/glycoPER ratio suggesting a strong shift away from OXPHOS activity towards aerobic glycolysis (Figure 2). Stimulation with both concentrations of PIC also increased PER levels at baseline and the %PER derived from glycolysis but this increase was significantly lower than that observed following LPS-stimulation (P<0.001). Further, the reduction in the mitoOCR/glycoP ER ratio was less pronounced suggesting that PIC-stimulated cells maintain higher levels of OXPHOS activity (Figure 2C). Despite similar basal PER levels, low and high concentrations of PIC differentially affected the ability of BMM to ramp up glycolysis following stress with Rot/AA. While cells stimulated with lower concentrations maintained their ability to increase glycolytic activity following exposure to Rot/AA, ce**l** s stimulated with higher concentrations were unable to do so, suggesting they may be functioning at their maximum glycolytic capacity (Figure 2A).

**Figure 2:**
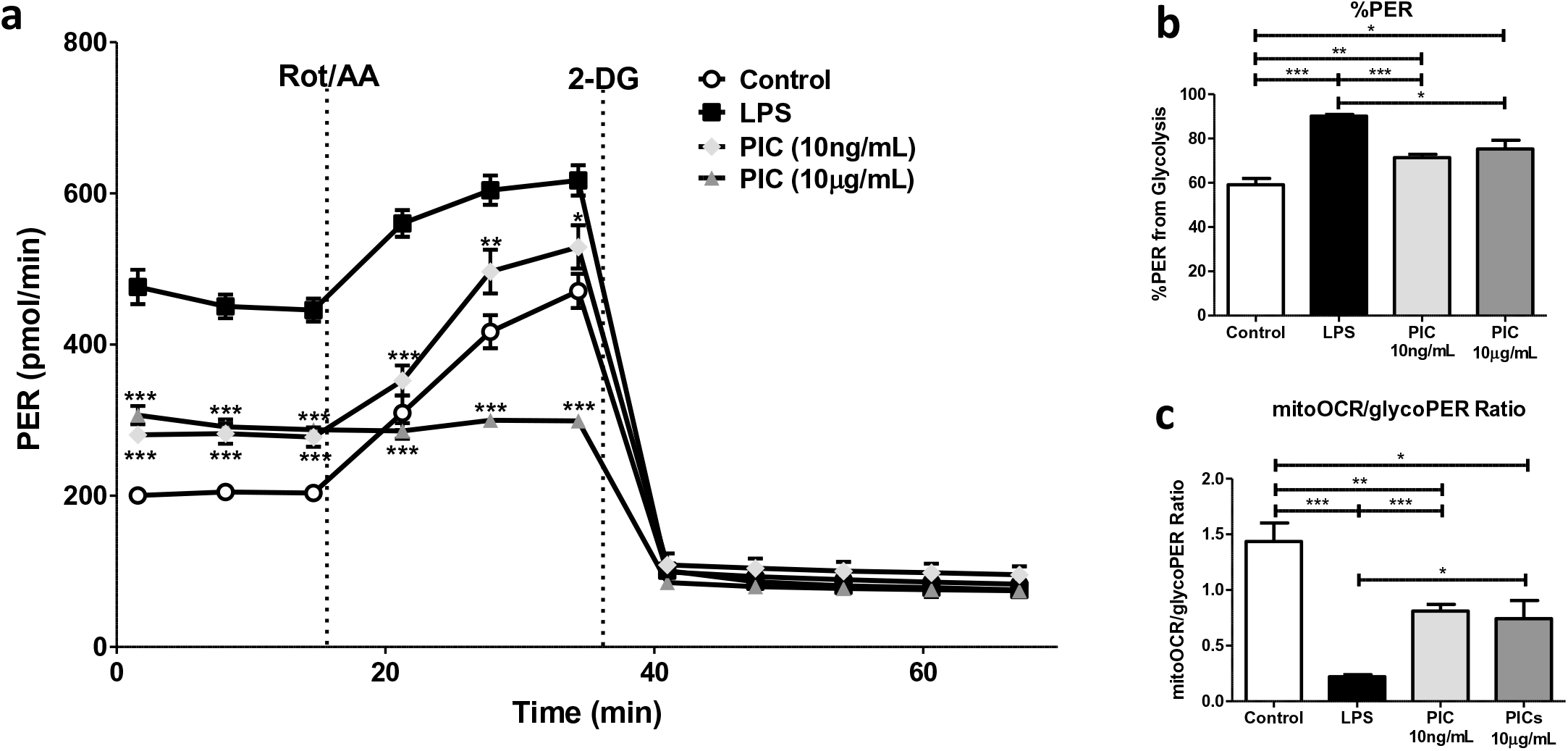
Macrophages activated using higher concentrations of Poly(I:C) are functioning near their maximum glycolytic capacity. BMDMs were seeded onto Seahorse XFp miniplates and treated with 100ng/mL LPS, 10ng/mL or 10μg/mL PIC for 18 hours. Glycolytic activit y, indicated by the proton efflux rate (PER) was measured using sequential injections of rotenone plus antimycin A (Rot/AA) and 2-deoxyglucose (2-DG) **(a)**, determining the %PER dependent on glycolysis **(b)** and the ratio of mitochondrial oxygen consumption rate (mitoOCR) to glycolyt ic PER **(c)**. Data represents mean ± SEM of four individual mice. The levels of significance shown in Figure 2a represent pairwise comparisons against LPS-treated macrophages (*p < 0.05, **p < 0.01, and ***p < 0.001).

### Maintenance of OXPHOS activity is an important feature of PIC but not LPS stimulation

To evaluate specific changes in OXPHOS activity, we used the Cell Mito Stress Test kit from Agilent. Features of OXPHOS activity were calculated based on changes in oxygen consumptio n rate (OCR) in response to successive injections of oligomycin (Oligo), carbonyl cyanide 4-(trifluoromethoxy)phenylhydrazone (FCCP) and ROT/AA. Consistent with the literature^9, 19, 36^, LPS stimulation dramatically reduced basal respiration, reduced mitochondrial ATP production and reduced the ability of cells to increase oxygen consumption (e.g. spare respiratory capacity [SRC]) following FCCP treatment (Figure 3). High concentrations of PIC also reduced levels of basal respiration, ATP production and SRC compared to untreated cells but this impairment was less severe than that observed for LPS (basal respiration p<0.05, SRC p<0.01, ATP production P=0.08). Interestingly, low concentrations of PIC did not alter basal respiration or ATP production but significantly reduced SRC suggesting these cells may have a reduced capacity to deal with stress^37^. Given these differences in glycolysis and oxygen consumption, the remaining experiments were performed using low concentration PIC (10ng/ml).

**Figure 3:**
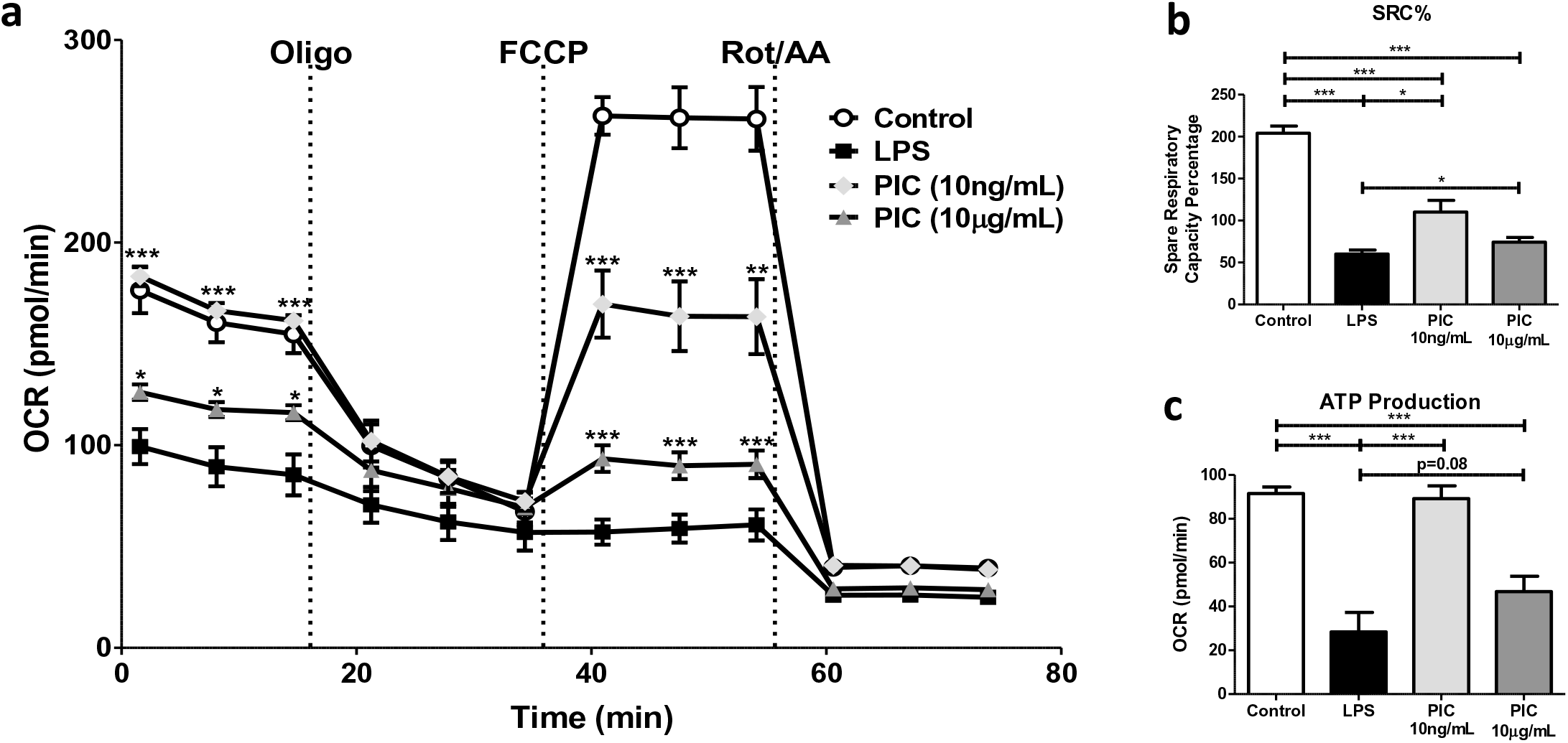
Poly(I:C) stimulation is linked to low sustained levels of oxidative phosphorylatio n (OXPHOS). Macrophages were plated onto Seahorse XFp miniplates and subsequently stimulated with 100ng/mL LPS, 10μg/mL or 10ng/mL PIC for 18 hours. OXPHOS function was assessed *via* successive Oligomycin (Oligo), Carbonyl cyanide-*p*-trifluoromethoxyphenylhydrazone (FCCP), and Rot/AA injections **(a)**, quantifying the spare respiratory capacity percentage (SRC%) **(b)** and ATP production **(c)**. Data represents mean ± SEM of four individual mice. The levels of significance shown in Figure 3a is based on pairwise comparisons to LPS-treated macrophages (*p < 0.05, **p < 0.01, and ***p < 0.001).

### Reduced glucose availability is associated with increased inflammatory and antiviral cytokine production in PIC-but not LPS-stimulated BMM

Macrophages are highly plastic cells whose responses are modified by environmental cues including nutrient availability^38–41^. A recent study in BMM showed that macrophages are less dependent on OXPHOS activity under condition of high glucose (>10 mM) and preferentially use aerobic glycolysis to rapidly produce ATP^23^. Given the differential ability of BMM to reprogram OXPHOS activity based on glucose availability, we next evaluated how glucose levels affected TLR3 and TLR4 signaling and cytokine production. Most studies have been performed in standard culture conditions, which represent supra-physiological concentrations of glucose. For these studies, BMM were stimulated with LPS (100ng/ml) and PIC (10ng/ml) in standard (25mM) and low glucose (0.5mM) conditions. For the remainder of this paper, standard culture conditions will be referred as high glucose. Glucose levels had no effect on pro-inflammatory cytokine production in LPS-stimulated cells (Figure 4) suggesting that even low glucose levels are sufficient to support TLR4 responses. Conversely, low glucose conditions increased pro-inflammatory (TNF-α, IL-6) and IFN-associated cytokine production (IFN-α, IFN-β, CXCL10) following PIC stimulation suggesting that glucose availability may fine tune the magnitude of the TLR3 response. Interestingly, we found that high levels of glucose increase baseline OCR levels, spare respiratory capacity and ATP production in untreated/resting ce**l** s. However, following PIC stimulation, high glucose further reduced basal OCR, spare respiratory capacity and ATP production suggesting these conditions may alter TLR3 associated mitochondrial reprogramming (Supplementary Figure S1).

**Figure 4:**
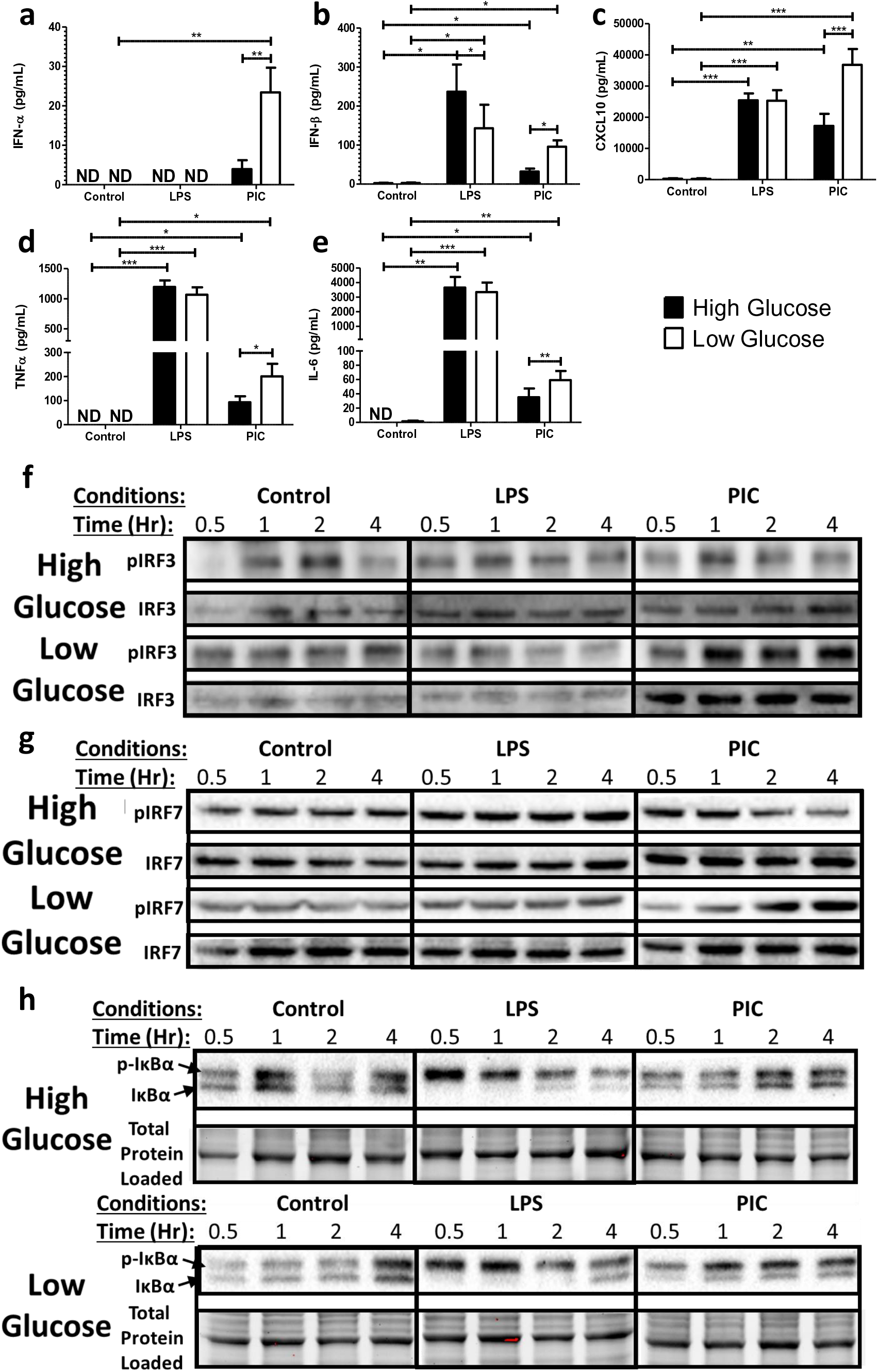
Low glucose conditions are associated with increased IRF activation and increased type I IFN production. Macrophages were stimulated with either 100ng/mL LPS or 10ng/mL PIC for 18 hours under high glucose (25mM) or low glucose (0.5mM) media conditions. Supernatant was collected for assessing antiviral (IFN-α, IFN-β, CXCL10) **(a-c)** and pro-inflammatory (TNF-α, IL-6) cytokine **(d-e)** expression. Cell lysates were harvested to quantify IRF3 **(f)**, IRF7 **(g)** and Iκbα **(h)** expression *via* immunoblotting. Data represents mean ± SEM of four individual mice (*p < 0.05, **p < 0.01, and ***p < 0.001).

To determine if this amplification was associated with altered signaling, we next evaluated alterations in TLR3 and TLR4 adaptor protein expression and transcription factor phosphoryla t io n following stimulation with LPS and PIC under high and low glucose conditions. While TRIF and TRAF6 expression was not affected by glucose levels (Supplementary Figure 2), low glucose was associated with a dramatic increase in total and phosphorylated IRF3 in PIC-stimulated BMM. Consistent with its more downstream role in TLR3 signaling^42, 43^, we also found IRF7 phosphorylation was delayed but sustained at high levels in low glucose conditions (Figure 4F and 4G). Furthermore, we found that levels of phosphorylated IκBα were increased in low glucose conditions in PIC but not LPS-stimulated cells, which may explain the increased TNF-α and IL-6 production following TLR3 engagement (Figure 4H). Collectively, these results suggest that high glucose may limit mitochondrial reprogramming and associated antiviral and pro-inflammatory signaling and cytokine production in a TLR3 specific manner.

### Reduced glucose availability is associated with altered mitochondrial membrane potential and ETC complex expression following TLR engagement

To better understand how glucose levels affect TLR3 associated alterations in OXPHOS activity, we examined alterations in mitochondrial membrane potential (MMP) and ETC complex expression in high and low glucose conditions. MMP was assessed using the fluorescent dye Tetramethylrhodamine (TMRM). As previously described^15^, LPS stimulation was associated with increased sequestration of TMRM by activated mitochondria (TMRM Mean Fluorescence Intensity [MFI]) in both high and low glucose conditions (Figure 5A). Alternatively, PIC-stimulation did not significantly increase levels of TMRM sequestration in positive cells. Instead, PIC was associated with a significant increase in the number of cells expressing low levels of TMRM, which further increased under low glucose conditions (24% vs. 38%, p=0.09). To evaluate if altered ETC flux contributes to altered membrane potential, we examined ETC complex expression following LPS and PIC stimula t io n in high versus low glucose conditions. In high glucose conditions, alterations in expression were highly variable across animals. LPS-stimulation moderately decreased complex II (SDHB) expression whereas PIC increased complex IV (COX4) (Figure 5B). Alterations in ETC complex expression were more pronounced in low glucose conditions. Specifically, both LPS and PIC were associated with decreased expression of complexes I and IV. The only alteration unique to PIC in the low glucose condition was the significant increase in complex III (Figure 5B). In addition to its role as a proton pump, complex III is a major generator of mtROS^44^ and may contribute to the amplification of inflammatory and antiviral cytokine production following TLR3 engagement.

**Figure 5:**
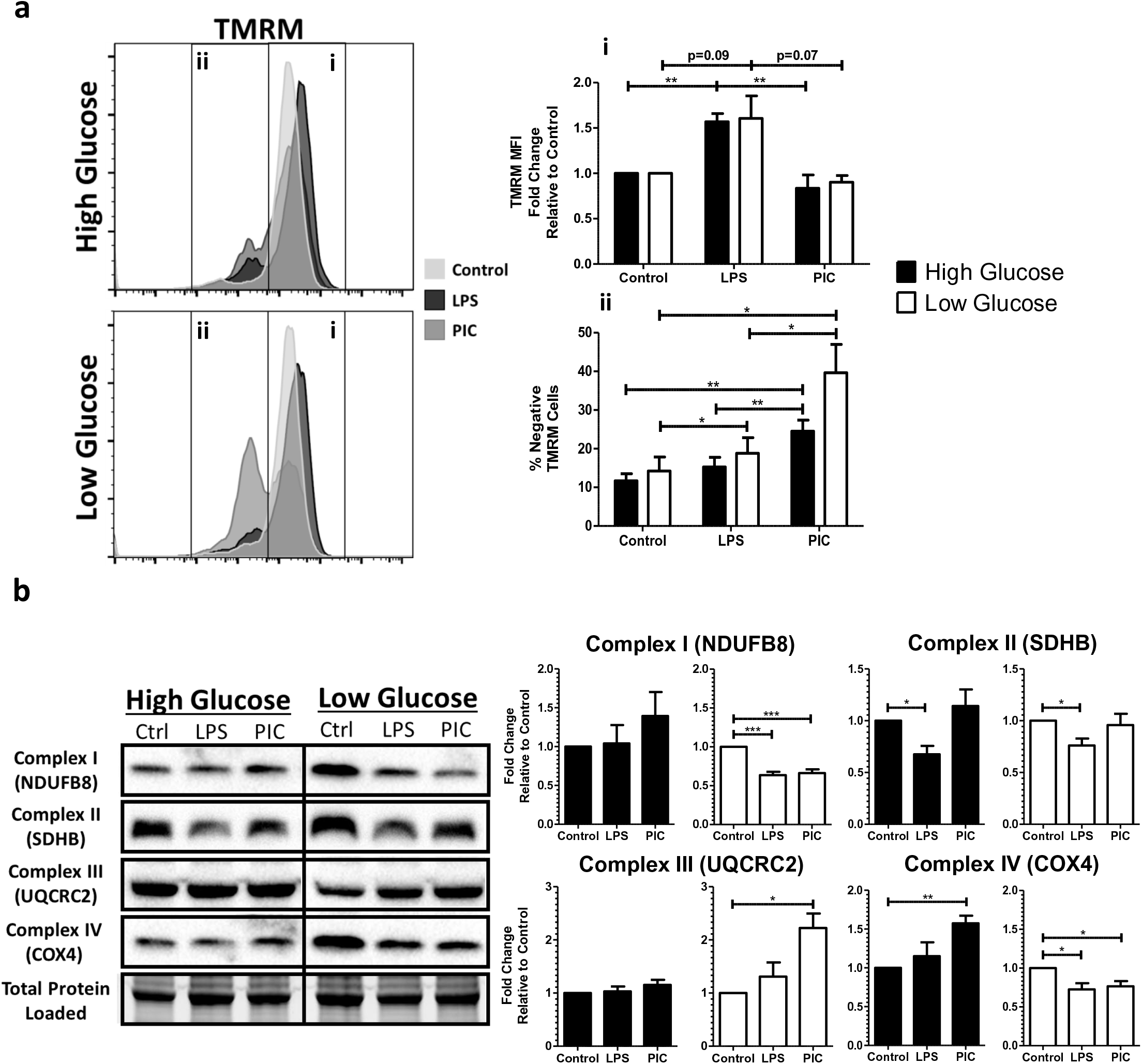
Poly(I:C) activation is linked to altered mitochondrial activity under low glucose conditions. BMDMs treated with LPS or PIC for 18 hours under high glucose or low glucose media conditions were characterized for differences in mitochondrial functio n. Tetramethylrhodamine (TMRM) staining was used to measure, *via* flow cytometry, mitochondr ia l membrane potential **(a)**. Core protein levels of Complexes I-IV of the electron transport chain was quantified *via* immunoblotting **(b)**. Data represents mean ± SEM of four individual mice (*p < 0.05, **p < 0.01, and ***p < 0.001).

### Flux through the ETC chain is required for inflammatory and antiviral cytokine production following TLR engagement

To evaluate the relative contribution of specific ETC complexes on cytokine production, BMM were stimulated with LPS (100ng/ml) and PIC (10ng/ml) under high and low glucose conditions in the presence or absence of Rotenone (Complex I inhibitor), Antimycin (Complex III inhibitor) and Cyanide (Complex IV inhibitor). Inhibition of Complex III and IV significantly reduced CXCL10 (III: ↓51%; IV: ↓72%) and TNF production (III: ↓54%; IV: ↓77%) following LPS stimulation (Figure 6A). Under low glucose conditions, inhibition of Complex I also limited LPS-associated cytokine production (CXCL10: ↓57%; TNF: ↓55%). Similarly, inhibition of Complexes I, III and IV significantly reduced inflammatory (TNF) and antiviral cytokine production (IFN-α, IFN-β, CXCL10) following PIC stimulation (Figure 6B, Supplementary Figure 3). The magnitude of this inhibition was further amplified under low glucose conditions, particularly for complex III (CXCL10: ↓70% vs ↓92%; TNF-α: ↓78% vs. ↓89%; IFN-β: ↓79% vs. ↓88%). These results suggest that ETC flux is required for inflammatory and antiviral cytokine production and that under low glucose conditions, alterations in complex III expression may play a central role in the amplification of these responses following TLR3 engagement.

**Figure 6:**
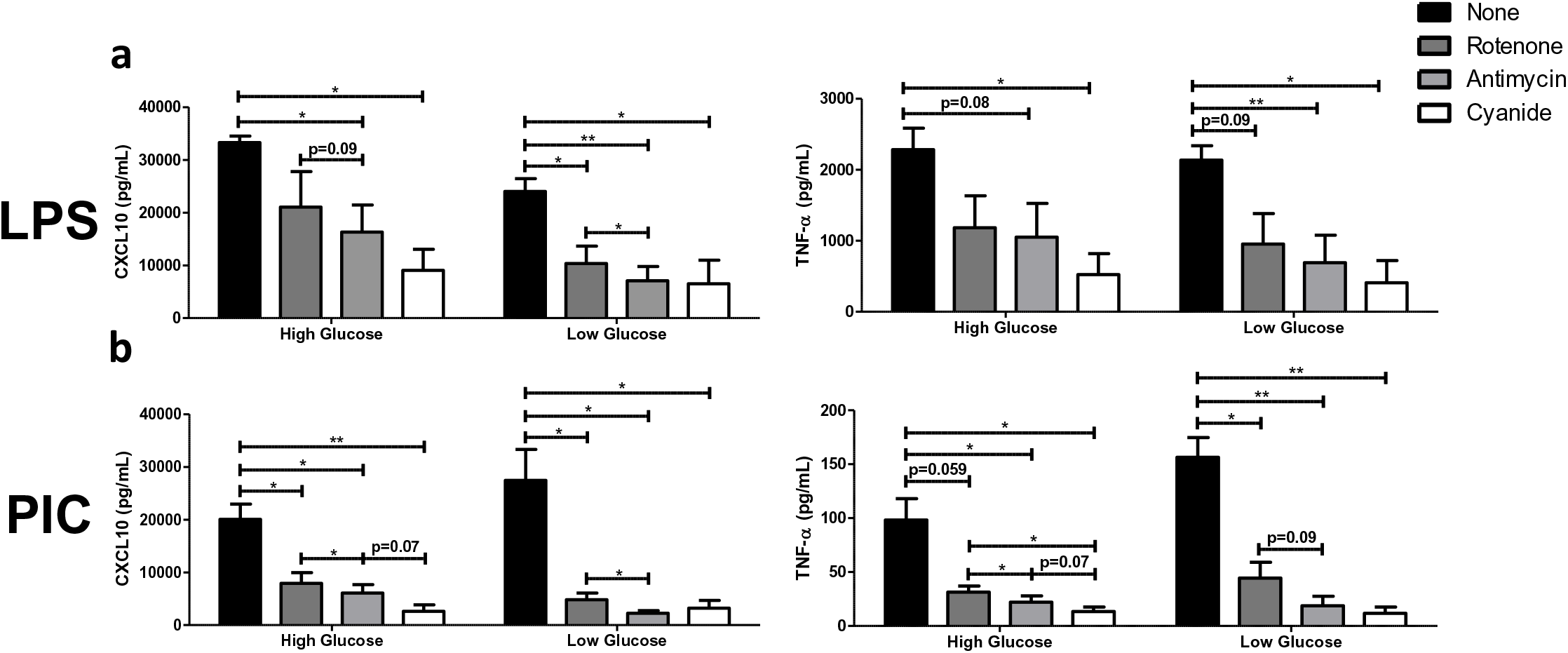
Targeting ETC activity reduces type I IFN-mediated responses during Poly(I:C) activation. LPS-**(a)** or PIC-**(b)** stimulated BMDMs were co-treated with a panel of ETC inhibitors (Rotenone, Antimycin, Cyanide) to assess the importance of mitochondrial function for antiviral responses. CXCL10 and TNF-α cytokine secretion was measured after 18 hours in high glucose or low glucose media conditions. Data represents mean ± SEM of three individual mice (*p < 0.05, **p < 0.01, and ***p < 0.001).

### Mitochondrial and cytosolic ROS accumulate in PIC stimulated BMM under low glucose conditions

Given the central role of complexes I and III in driving mitochondrial ROS production, we next quantified mitochondrial superoxide production using the fluorescent probe MitoSOX^TM^ Red. As previously reported^45^, LPS stimulation was associated with increased mitochondr ia l superoxide production compared to untreated BMM. This increase was unaffected by glucose availability (Figure 7A). PIC stimulation was also associated with increased mitochondr ia l superoxide production (Figure 7A); however, its production was further increased under low glucose conditions. To determine if this superoxide accumulation was associated with altered antioxidant expression, cellular levels of superoxide dismutase 2 (SOD2) and mitochondr ia l glutathione peroxidase 4 (mtGPX4) were evaluated *via* western blots. In high glucose conditions, both LPS and PIC stimulated cells significantly increased SOD2 (LPS-FC=2.56±0.44; PIC-FC=1.75±0.40) and mtGPX4 expression (LPS-FC=1.40±0.07; PIC-FC=1.48±0.22) levels (Figure 7B). Alternatively, while LPS upregulated both antioxidant proteins under low glucose conditions (SOD2-FC=2.11±0.40; mtGPX4-FC=2.09±0.41), levels of SOD2 (FC=0.83±0.22) and mtGPX4 (FC=1.35±0.38) were not altered following PIC. This may contribute to the accumulation of superoxide in the mitochondria. To evaluate if these alterations affect the accumulation of cytosolic ROS, we used CellROX^TM^ Orange and quantified H2O2 levels in cell lysates. CellROX^TM^ Orange has a high affinity for hydroxyl radicals, H2O2 and superoxide. BMM treated with LPS exhibit increased levels of cytosolic ROS in both high and low glucose conditions (Figure 7C). Conversely, increased cytosolic ROS was only observed in low glucose conditions following PIC stimulation (Figure 7C and 7D). These results suggest that low glucose conditions are associated with increased mitochondrial and cytosolic ROS accumulation following PIC stimulation, which may contribute to the amplification of the cytokine response.

**Figure 7:**
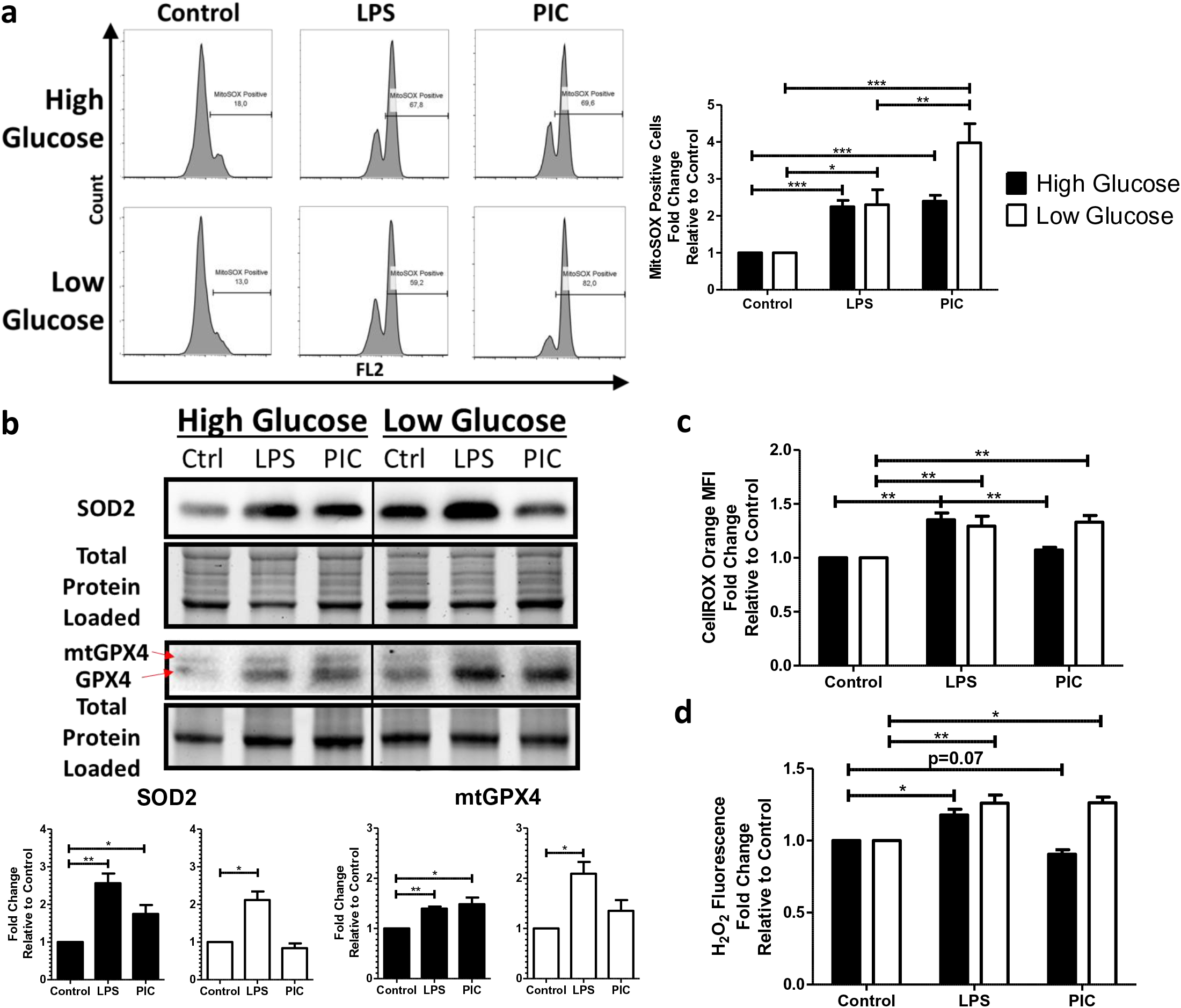
Poly(I:C) activation promotes mitochondrial ROS production and accumulation. Macrophages treated either with LPS or PIC for 18 hours under high glucose or low glucose media conditions were examined for differences in redox metabolism. Mitochondrial superoxide production was measured using MitoSOX Red^TM^ **(a)**. Protein levels of antioxidant proteins superoxide dismutase (SOD2) and mitochondrial glutathione peroxidase 4 (mtGPX4) were measured via immunoblotting **(b)**. Cytosolic ROS production was measured using CellROX Orange **(c)**. Hydrogen peroxide levels were quantified using the Cell-based Hydrogen Peroxide Assay kit **(d)**. Data represents mean ± SEM of four individual mice (*p < 0.05, **p < 0.01, and ***p < 0.001).

### Mitochondrial and cellular ROS play a central role in TLR3 and TLR4 associated cytokine production

To evaluate if ROS contributes to cytokine production following TLR3 and TLR4 engagement, ce**l** s were stimulated with either LPS (100ng/ml) or PIC (10ng/ml) under high and low glucose conditions in the presence or absence of antioxidants (MitoTEMPO, *N*-acetylcysteine) or an inhibitor of superoxide production (S3QEL). MitoTEMPO (MT) is a mitochondria-spec if ic antioxidant that selectively scavenges mitochondrial superoxide. Alternatively, *N*-acetylcysteine (NAC) boosts glutathione synthesis reducing total overall cellular ROS production. S3QEL selectively inhibits superoxide production from the outer Q-binding site of the ETC complex III without altering OXPHOS. Inhibition of mitochondrial ROS had differential effects on cytokine production following LPS stimulation (Figure 8A; Supplementary Figure 4). Whereas TNF-α (MT: ↓50%; S3QEL: ↓48%) and IFN-β levels (MT: ↓60%; S3QEL: ↓61%) were decreased following MT and S3QEL production, CXCL10 levels were unaffected. CXCL10 production was only reduced following NAC treatment (CXCL10: ↓71%) suggesting cytosolic ROS may play a more important role in its production following LPS stimulation. Alternatively, inhibition of mitochondrial ROS significantly reduced TNF-α, CXCL10, IFN-α and IFN-β production in PIC-activated macrophages (Figure 8B; Supplementary Figure 4). The magnitude of S3QEL inhibit io n was further amplified under low glucose conditions (TNF-α: ↓54% vs. ↓66%; CXCL10: not significant vs ↓66%, IFN-β: ↓61% vs. ↓73%) suggesting complex III plays an important role in the amplification of the TLR3 responses. Similar to LPS, NAC treatment had the most pronounced effects on cytokine production in both and high low glucose conditions suggesting that both mitochondrial and cytosolic ROS contribute to TLR3 mediate cytokine production.

**Figure 8:**
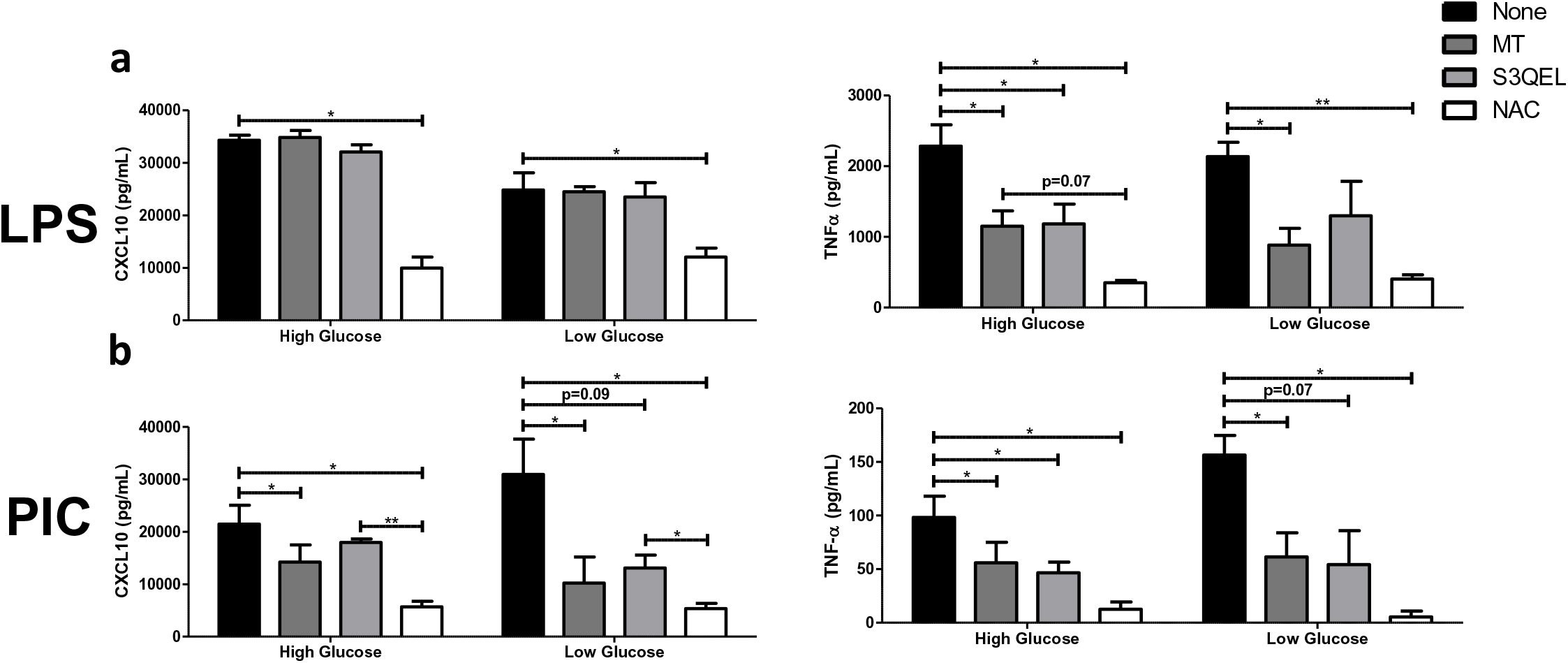
Type I IFN production can be inhibited by altering mtROS generation during Poly(I:C) activation. LPS-**(a)** or PIC-**(b)** stimulated BMDMs were co-treated with a panel of mtROS (MT, S3QEL, NAC) modulators to assess the importance of mitochondrial function for antiviral responses. CXCL10 and TNF-α cytokine secretion was measured after 18 hours in high glucose or low glucose media conditions. Data represents mean ± SEM of three individual mice (*p < 0.05, **p < 0.01, and ***p < 0.001).

**Figure 9:** Proposed model of the relationship between mitochondrial ROS and IFN-mediate d responses in Poly(I:C)-stimulated macrophages. Mitochondrial-derived ROS produced during Poly(I:C) activation leads to increased IRF3/7 activation and subsequent type I IFNs and CXCL10 expression. This effect is amplified under low glucose, with increased Complex III expression and reduced mitochondrial membrane potential. Targeting either electron flow through Complex III or mitochondrial superoxide production leads to reduced IRF3/7 activation as we**l** as type I IFNs and CXCL10 expression.

## Discussion

Increasing evidence suggests that mitochondria play a critical role in driving innate immune responses against bacteria and viruses^15, 16, 20–23, 33^. However, it is unclear if specific features of mitochondrial reprogramming contribute to pathogen specific immune responses or how nutrient availability may affect these processes. In the current study, we found that TLR3 and TLR4 engagement uniquely remodeled ETC complex expression, resulting in differential accumula t io n of mitochondrial and cytosolic ROS. This differential ROS production is required to support ligand specific inflammatory and antiviral cytokine profiles. We also found that the magnitude of TLR3 but not TLR4 responses were modulated by glucose availability. Under conditions of low glucose, TLR3 engagement was associated with increased ETC complex III expression, increased mitochondrial and cytosolic ROS and increased inflammatory and antiviral cytokine production. This increased cytokine production was selectively reversed by targeting superoxide production from the outer Q-binding site of the ETC complex III. Collectively, these findings suggest that the ETC may act as a selective rheostat of macrophage function that regulates not only the nature (antibacterial vs. antiviral) but the magnitude of the response, which may depend on nutrient availability.

It is widely accepted that inflammatory macrophages undergo metabolic reprogramming to support cytokine production and effector functions. In LPS-stimulated cells, reprogramming is associated with a near complete inhibition of OXPHOS and an increased reliance on aerobic glycolysis to support rapid energy production^9, 15, 17, 36, 46^. This switch is driven by altered flux through the tricarboxylic acid (TCA), which repurposes mitochondrial function to support superoxide production and drive intracellular anti-bacterial responses^15, 17^. While it was initially assumed all “inflammatory” stimuli induce similar responses, increasing evidence suggests this may not be the case^22, 23, 47^. In the current study, we found that PIC stimulation inhibited OXPHOS activity in a dose dependant manner. However, even at its highest concentrations (10μg/ml), PIC did not completely inhibit OXPHOS activity and some level of cellular respiration was maintained. Consistent with these findings, Yoshizumi et al. found that RLR-mediated responses in macrophages are dependent on OXPHOS both in vitro and in vivo. In BMM, disruption of cellular respiration severely impaired RLR induced interferon and proinflammatory cytokine production^23^. In mice, inhibition of OXPHOS was found to increase susceptibility to viral infection and induce significant inflammation in the lung^23^. Wu et al. found that TLR9 engagement and type I IFN production in plasmacytoid dendritic cells was associated with increased OXPHOS activity. This increase was fueled by fatty acid oxidation (FAO) and was required for full cellular activation^48^. In vivo, inhibition of FAO resulted in a diminished capacity to control lymphocytic choriomeningitis virus^48^. Several IFN-stimulated genes, such as ISG15, have also been linked to the regulation of mitochondrial function during viral infection suggesting a secondary wave of mitochondrial reprogramming may occur following the TLR engagement and the induction of type I IFN responses^49^. Collectively, these studies suggest that some level of OXPHOS activity may be required to mount functional antiviral immune responses but that these responses may vary by ligand and cell type.

In addition to altered cellular respiration, LPS and PIC stimulation were associated with alterations in mitochondrial membrane potential (MMP). MMP is generated by the proton pumps of the ETC (Complexes I, III and IV) to support mitochondrial ATP production^50^. Various studies have reported altered MMP following macrophage activation. Mills et al. found that LPS stimula t io n was associated with increased MMP *via* reverse electron flow (RET). This RET was required to drive electrons back towards Complex I in order to support mitochondrial ROS production and antimicrobial effector functions^15^. Koshiba et al, have found MMP is required for MAVS-mediated antiviral signaling. Specifically, they found that inhibition of mitochondrial fusion resulted in a widespread loss of MMP. This loss in MMP correlated with the level of inhibition of RLR-induced antiviral responses^22^. Unlike LPS, we found that PIC stimulation was associated with decreased MMP in a subset of cells. While it is unclear what exactly these cells represent, Tal et al. reported that when autophagy is inhibited, increased accumulation of dysfunctional mitochondria results in increased mitochondrial ROS production, which drives excess RLR signaling^26^. Further studies are required to determine if this subset of TMRM low cells are the main producers of ROS in our model system.

Recent studies have provided evidence that changes in the ETC, particularly in complex I and II, contribute to the regulation of antibacterial immune responses^15, 33^ however, it unclear is if similar remodeling occurs during antiviral responses. Here, we found that PIC stimulated cells undergo differential remodeling of the ETC, particularly with limited glucose availability. Standard cell culture conditions represent supra-physiological levels of glucose (25mM vs. 5-7mM in fasting blood from non-diabetic individuals^51^) and may alter mitochondrial reprogramming *in vitro*^52^. Under low glucose conditions, both LPS and PIC were found to downregulate complex I and IV expression. Interestingly, only PIC stimulation also increased complex III expression. This increased expression was associated with the amplification of the TLR3 cytokine production, which was reversed by the selective inhibition of ROS production by complex III. In support of our results, others have linked complex III to immune activation and function. In T cells, Sena et al. demonstrated that specific deletion of Rieske iron-sulfur protein (RISP), an essential component of Complex III, reduced mtROS production, nuclear factor of activated T cells (NFAT) activatio n, IL-2-mediated T cell activation and antigen-specific expansion in vivo^53^. Alternatively, ablation of complex III in regulatory T cells has been shown to reduce their inhibitory capacity without altering cell proliferation and survival^54^. In macrophages, listeria infection has been shown to increase ROS production *via* complex III, which drives NF-kappa-B essential modulator (NEMO) dimerization, increasing inhibitor of nuclear factor kappa-B kinase (IKK) activation, NF-κB signaling and cytokine production^47^ Our study is among the first to identify associations between complex III mediated ROS production and TLR3 antiviral immune responses.

While ROS are generally considered toxic and damaging, increasing evidence suggests they also influence cellular signaling^55, 56^. Superoxide and its more stable derivative hydrogen peroxide have been shown to regulate a variety of biological responses such as cell proliferation, differentiat io n, and migration^55^. In the current study, we found that both mitochondrial superoxide and cytosolic ROS contribute to inflammatory and antiviral cytokine production following TLR engagement and that differential accumulation of ROS across these compartments may contribute to pathogen specific responses. Furthermore, under low glucose conditions, we found PIC but not LPS was associated with increased mitochondrial superoxide and cytosolic ROS production, which consequentially amplified cytokine production in these cells. These results suggest that the dynamic regulation of ROS production, likely through the modulation of complex III, may act as a rheostat that regulates the magnitude of antiviral immune responses. Consistent with these findings, Agod et al. found that mitochondrial superoxide drives increased MAVS protein expression in plasmacytoid DCs, increasing Akt and IRF3 activation and subsequent type I IFN production^57^. Wang et al. showed shRNA knockout of SOD2 in cell lines increased viral replication and reduced antiviral responses^58^, likely a result of decreased mitochondrial H_2_O_2_ production, a known redox-sensitive activator of NF-κB and IRF signalling^47, 59–61^. While we believe that mitochondrial superoxide and associated hydrogen peroxide production are the main drivers of cytokine production in our system, we cannot exclude the possibility that alternative cytosolic sources of ROS may also contribute. NADPH oxidase (NOX)-generated ROS during respiratory syncytial virus (RSV) and herpes simplex virus (HSV) infections can activate both NF-κB and IRF signalling^60, 61^. Similarly, Yang et al. reported that high concentrations of PIC increase NOX2 activity and ROS production in BMMs, which was required for signal transduction and activator of transcription 1 (STAT1)-mediated signalling^62^. A similar phenomenon was observed by To et al. using TLR7 ligands^63^. Further studies are required to determine the specific contribution of mitochondrial vs. cytosolic derived ROS in driving these processes and the distinct roles of superoxide vs. hydrogen peroxide on signaling and effector function.

Taken together, our results suggest that dynamic remodeling of the ETC complex expression represents a mechanism by which macrophages modulate cytokine production following TLR engagement. We found that this remodeling was associated with differential accumulation of mitochondrial vs. cytosolic ROS, which may drive ligand specific cytokine profiles. We hypothesize that this differential accumulation may be driven by dependence on complex I (LPS) versus complex III (PIC) for ROS production. Specifically, that LPS associated ROS production depend on RET whereas PIC depends on the maintenance of low levels of OXPHOS activity. Interestingly, we also found that TLR3 but not TLR4 associated mitochondrial reprogramming was dependent on glucose availability in the microenvironment. Supra-physiological levels of glucose have been shown to decrease a cell’s dependence on mitochondria for energy production^52, 64^. Similarly, our results suggest high glucose conditions may also alter mitochondr ia l reprogramming associated with TLR3 engagement. Accordingly, it is important to develop a detailed understanding of these processes in a variety of bacterial and viral infections to identify new therapeutic approaches to help boost specific and functional effector functions.

## Methods

### Reagents

Lipopolysaccharide (LPS) and high molecular weight Poly(I:C) (PIC) were purchased from InvivoGen. MitoTEMPO (MT), *N*-acetylcysteine (NAC), antimycin A (AA), rotenone (ROT), potassium cyanide, 2-deoxyglucose (2-DG), oligomycin (OM) and carbonyl cyanide-*p*-trifluoromethoxyphenylhydrazone (FCCP) were acquired from Sigma-Aldrich while S3QEL-2 was purchased from Cedarlane. IL-1β, IL-6, IL-10, TNF-α, and CXCL10 ELISA kits were purchased from R&D Systems. The IFN-α/IFN-β 2-Plex Mouse ProcartaPlex^TM^ Luminex Panel kit used was from Invitrogen. Tetramethylrhodamine, methyl ester (TMRM), MitoSox Red and CellROX Orange probes were from ThermoFisher. All antibodies used in this study can be found in Table S1. Antibodies against Complex II (SDHB) was from Abcam while antibodies recognizing SOD2 was purchased from Cell Signalling Technology. Antibodies targeting IRF3, pIRF3 (Ser385), IRF7, pIRF7 (Ser477), GPX4, Iκbα, Complexes I (NDUFB8), III (UQCRC2) and IV (COX4) were purchased from ThermoFisher.

### BMM culturing and stimulation

Total bone marrow cells were collected from the tibias and femurs of 6-13-week-old C57BL/6 mice, cryopreserved in a 90% FBS/10% DMSO solution, and frozen until use. Ce**l** s were cultured for ten days in DMEM media with 10% fetal bovine serum, 1% penicillin/streptomycin (Life Technologies), and 15% L929 fibroblast cell-conditioned medium on a 100mm Petri dish as previously described^65^. On day 10, differentiated bone marrow-derived macrophages (BMM) were detached, counted and plated into tissue-culture treated plates at 1×10^6^ ce**l** s/mL. BMMs were stimulated with 100ng/mL LPS, 10ng/mL or 10μg/mL PIC under high (DMEM medium supplemented with 25mM glucose) or low glucose conditions (DMEM medium supplemented with 0.5mM glucose). The relative contribution of the ETC and ROS production on BMM inflammatory and antiviral cytokine production were assessed by co-treating stimulated cells with 1μM ROT, 5μM AA, 5mM cyanide, 500μM MT, 5mM NAC, or 5μM S3QEL-2.

### Cytokine Quantification

After the 18-hour stimulation, cytokine production was assessed in culture supernatants. IL-1β, IL-6, IL-10, TNF-α, and CXCL10 levels were assessed by ELISAs according to the manufactures instructions (R&D Systems). IFN-α and IFN-β levels were measured using IFN-α/IFN-β 2-Plex Mouse ProcartaPlex^TM^ Luminex Panel (Invitrogen).

### Western Blot Analysis

Untreated and stimulated BMMs (1×10^6^ cells) were lysed directly in the cell culture vessel using Pierce RIPA buffer (ThermoFisher) supplemented with HALT^TM^ Protease and Phosphatase Inhibitor (ThermoFisher). Total protein was quantified using the DC assay (Bio-Rad.) and resolved on a TGX^TM^ FastCast^TM^ Acrylamide gels (Bio-Rad). Gels were imaged directly using the Stain-Free application of a ChemiDoc XR (Bio-Rad) prior to transferring onto a PVDF membrane. Membranes were blocked overnight in 5% non-fat dry milk (w/v), washed and incubated overnight with the appropriate primary antibody. Horseradish peroxidase-conjugated secondary antibodies and Clarity^TM^ Western ECL Blotting Substrate (Bio-Rad) were used to visualize specified protein bands. Protein densitometry was analyzed according to previously described methodology^66^.

### Assessment of Mitochondrial Function by Flow Cytometry

BMMs were plated on 100mm Petri dishes and stimulated with 100ng/mL LPS or 10ng/mL and 10μg/mL PIC for 18 hours. Ce**l** s were then washed and stained with fluorescent probes according to the manufacturer’s instructions (30mins treatment at 37°C in select solutions). Mitochondr ia l membrane potential was measured using 10nM TMRM. Mitochondrial and Cellular ROS were monitored using 2.5μM MitoSOX Red in PBS and 5μM CellROX Orange in normal high glucose media, respectively. Cellular levels of fluorescence were quantified using an Attune NxT Flow Cytometer (ThermoFisher) and the results were analyzed using FlowJo Software. Results are reported as the percentage of positive cells and as mean fluorescence intensity (MFI), the latter being used to describe the level of expression on a population of positive cells.

### Quantification of cellular hydrogen peroxide production

BMMs were plated onto 96-we**l** black plates at 50,000 ce**l** s/well and stimulated with 100ng/mL LPS or 10ng/mL PIC under high or low glucose conditions for 1 hour. Ce**l** s were then washed before using the Cell-based Hydrogen Peroxide Assay Kit (Abcam) to measure H2O2 production. Cells were incubated with the AbGreen H2O2 indicator for 30 minutes before monitoring the relative difference in fluorescence using a fluorescence microplate reader (490nm Ex/520nm Em).

### Metabolic Extracellular Flux Analysis

BMMs were plated onto Seahorse XFp cell culture miniplates at 50,000 ce**l** s/well (Seahorse Bioscience) and stimulated with 100ng/mL LPS or 10ng/mL and 10μg/mL PIC for 18 hours. Extracellular acidification rate (ECAR) and oxygen consumption rate (OCR) were evaluated using a XFp Flux Analyzer (Seahorse Bioscience). Baseline ECAR and changes in glycolytic rate were assessed using the Seahorse XFp Glycolytic Rate Assay Kit (Agilent) according to the manufacturer’s instructions. Basal respiration, ATP production-coupled respiration, maximal and reserve capacities and non-mitochondrial respiration were assessed using the Seahorse XFp Cell Mito Stress Test Kit (Agilent).

### Statistical Analyses

Data used in this study was analyzed using GraphPad Prism software. Values shown represent the mean ± SEM of biological replicates, where the number of replicates are reported in the figure legends. Statistical significance was calculated using a paired Student’s *t*-test (*p < 0.05, **p < 0.01, and ***p < 0.001).

## Acknowledgements

Funding provided by a research development grant provided by the Carleton University Research Office and NSERC (RGPIN-2019-06214).

## Author Contributions

A.J., A.E. and A.A. isolated the bone marrow progenitor cells from mice. D.A., A.K., A.G. and E.C. designed the experiments. D.A. conducted the experiments. D.A., D.R. A.K., A.G. and E.C. analyzed the data and interpreted all the results. D.A., D.R., A.J., A.E., A.A., A.K., A.G. and E.C. all contributed to the writing and revising of this manuscript.

## Supplementary Information

**Supplementary Table S1:**
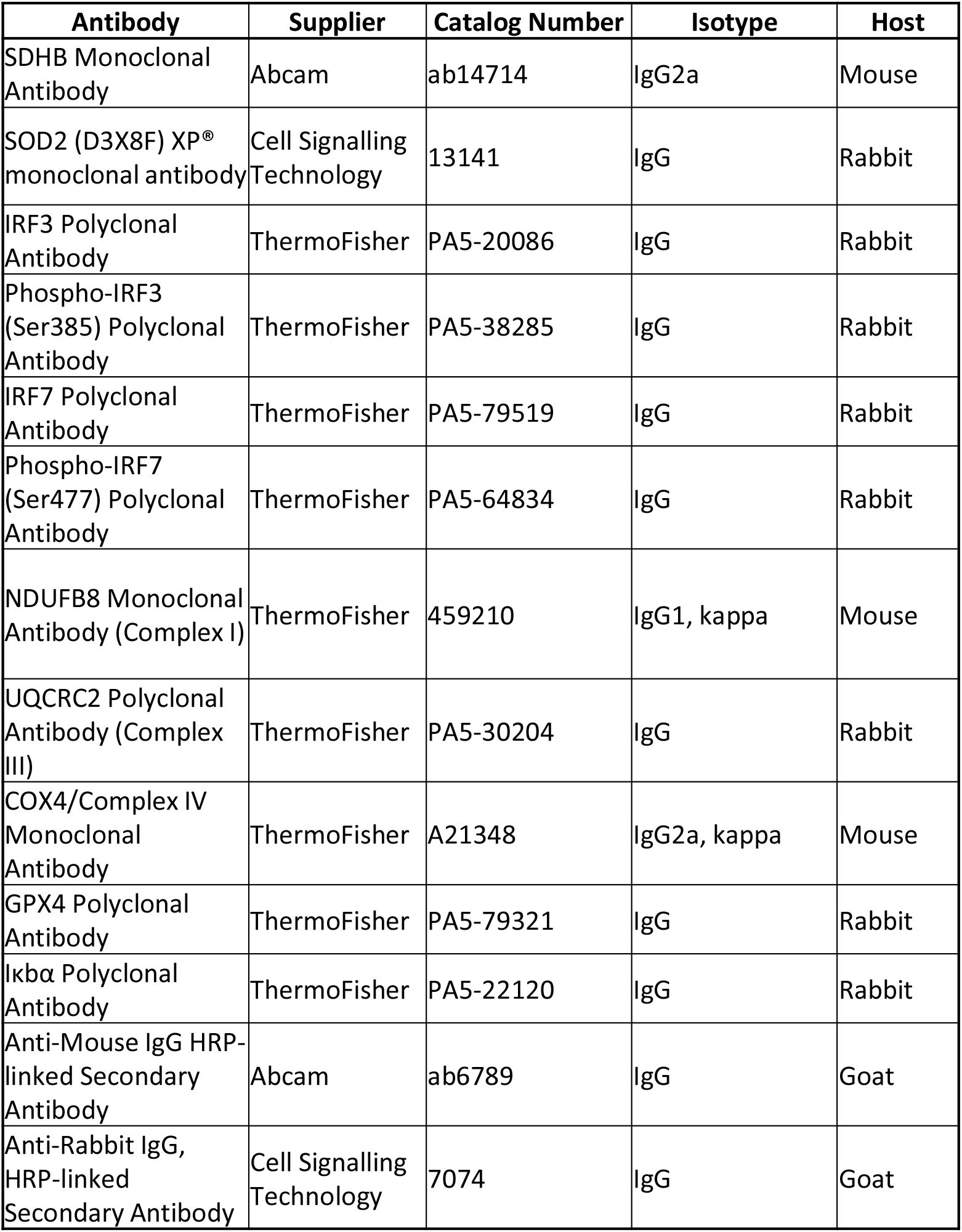
List of antibodies used in this study, referenced from Methods.

**Supplementary Figure S1:**
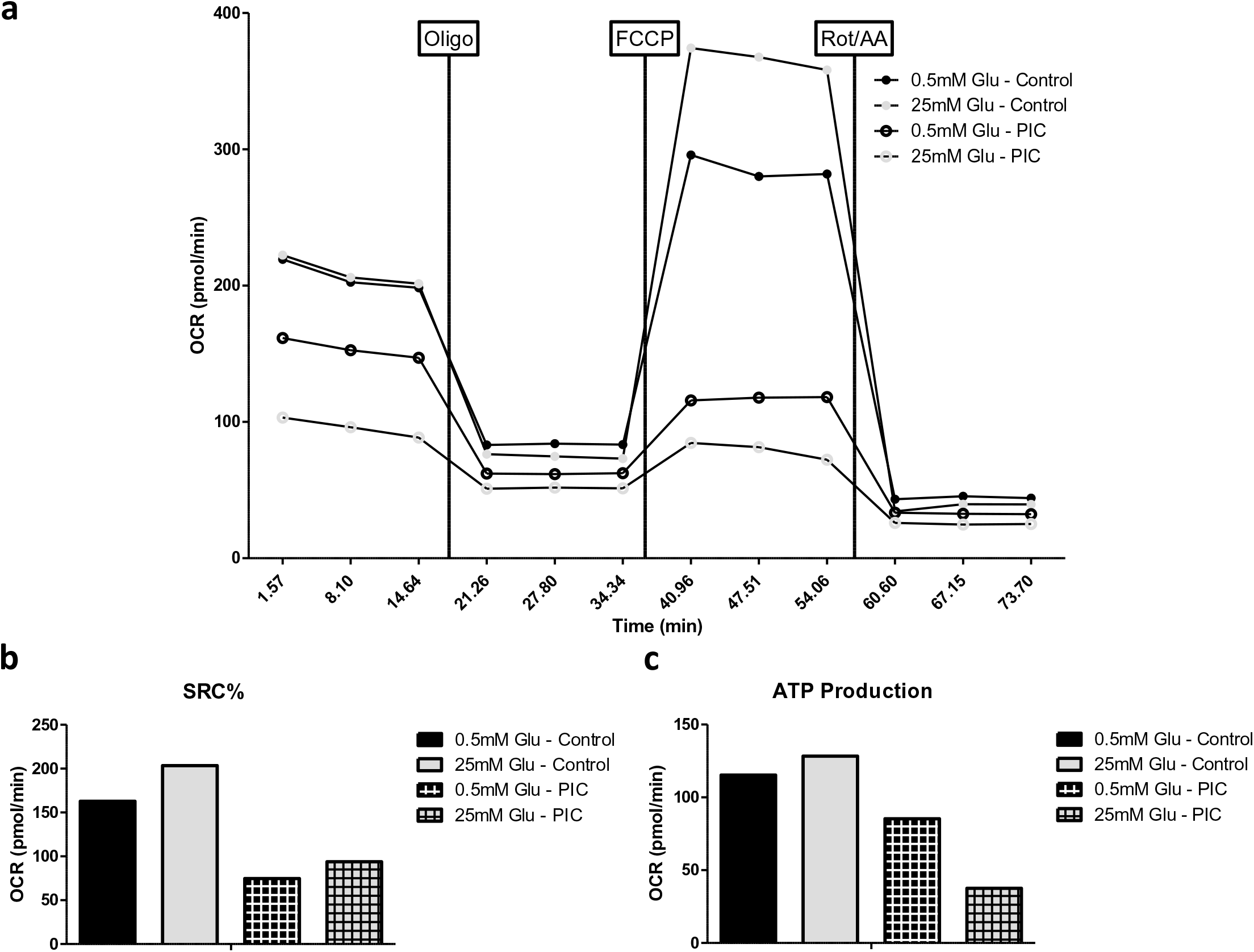
PIC activation is associated with increased OXPHOS function under low glucose conditions. BMDMs were plated onto Seahorse XFp miniplates and treated with 10μg/mL PIC for 18 hours. OXPHOS function was assessed *via* successive Oligomycin (Oligo), Carbonyl cyanide-*p*-trifluoromethoxyphenylhydrazone (FCCP), and Rot/AA injectio ns **(a)**. Quantification of the spare respiratory capacity percentage (SRC%) **(b)** and ATP production **(c)**. Data shown represents a test run using one representative animal.

**Supplementary Figure S2:**
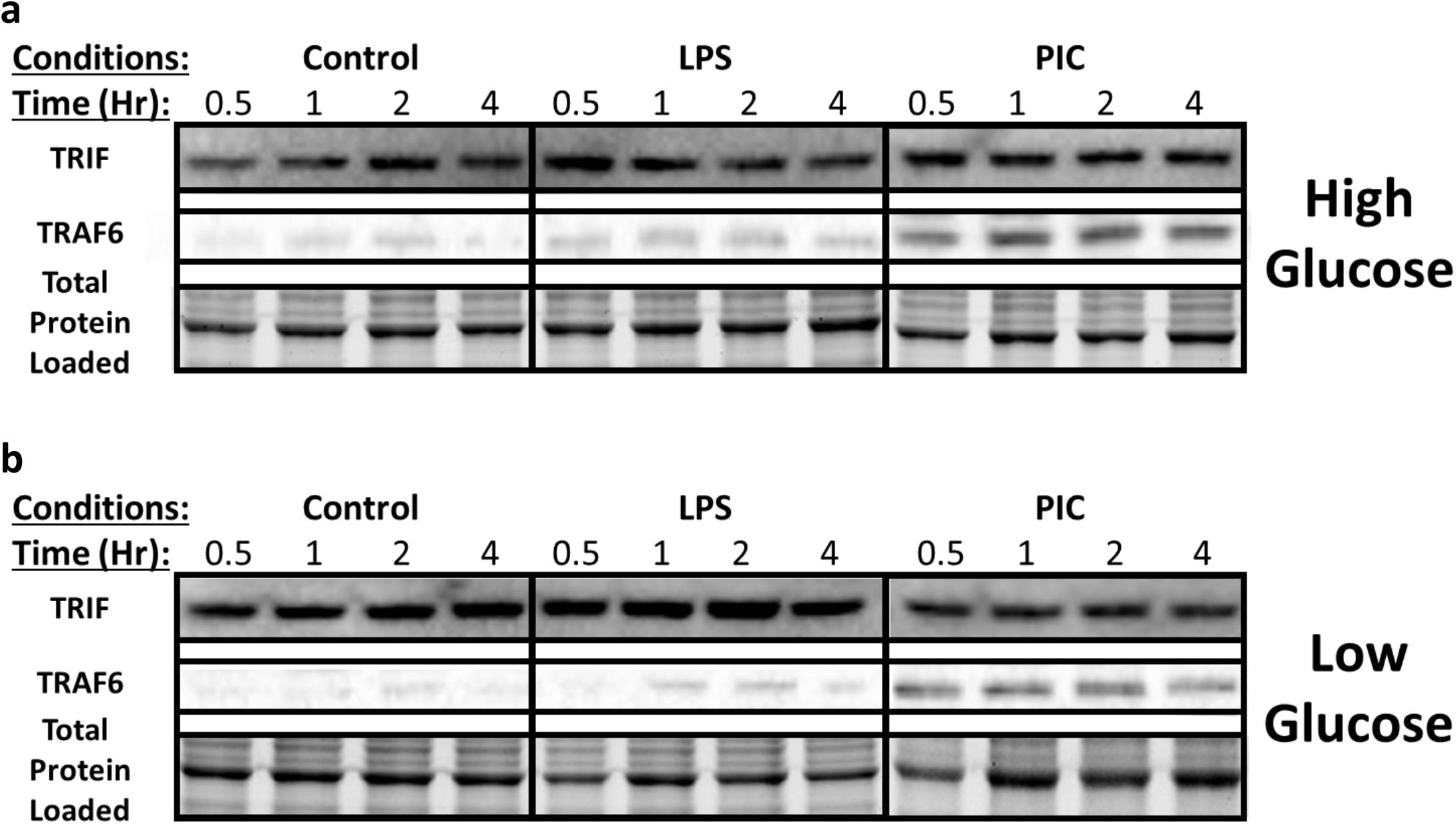
TRIF and TRAF6 expression during PIC activation is not affected by glucose levels. Macrophages stimulated either with LPS or PIC for 18 hours under high glucose **(a)** or low glucose **(b)** media conditions were examined for differences in TLR signaling. Protein levels of TRIF and TRAF6 were measured via immunoblotting. Data shown represents blots from a representative animal.

**Supplementary Figure S3:**
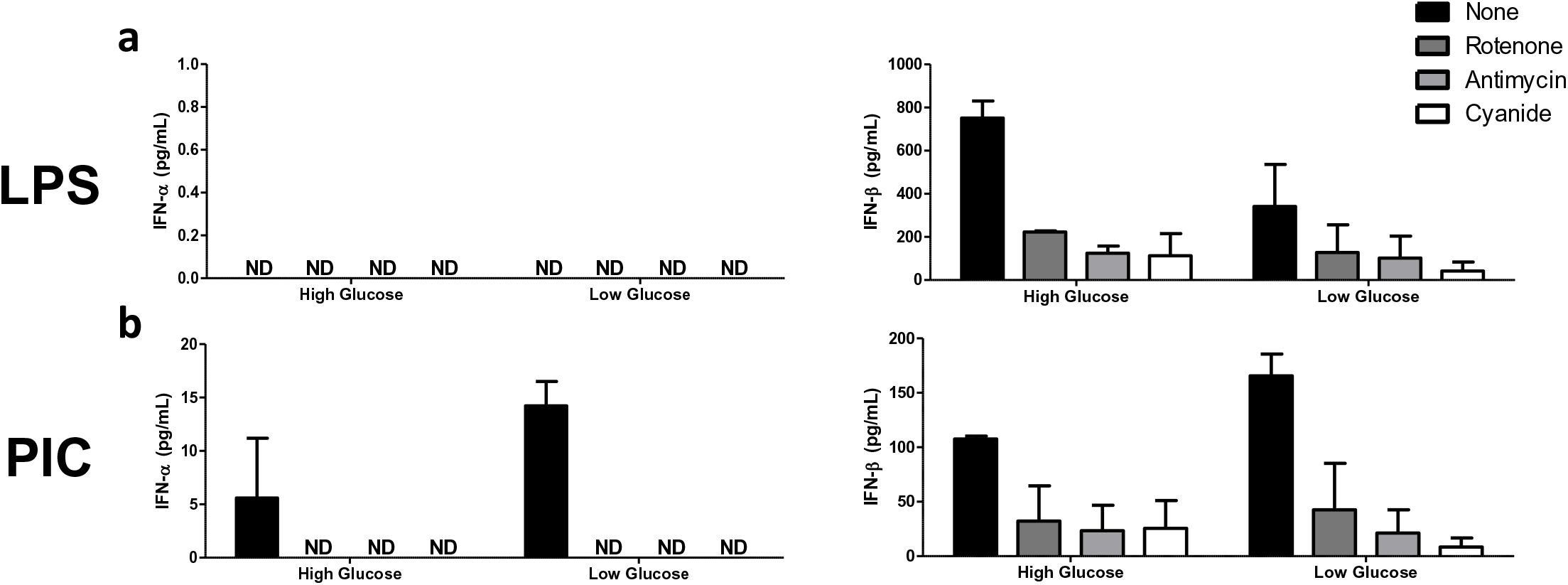
Targeting ETC function leads to complete loss of IFN responses. LPS-**(a)** or PIC-**(b)** stimulated BMDMs were co-treated with a panel of ETC (Rotenone, Antimycin, Cyanide) inhibitors to assess the importance of mitochondrial function to antivira l responses. CXCL10 and TNF-α cytokine secretion was measured after 18 hours in high glucose or low glucose media conditions. Data shown represents a mean of two individual animals.

**Supplementary Figure S4:**
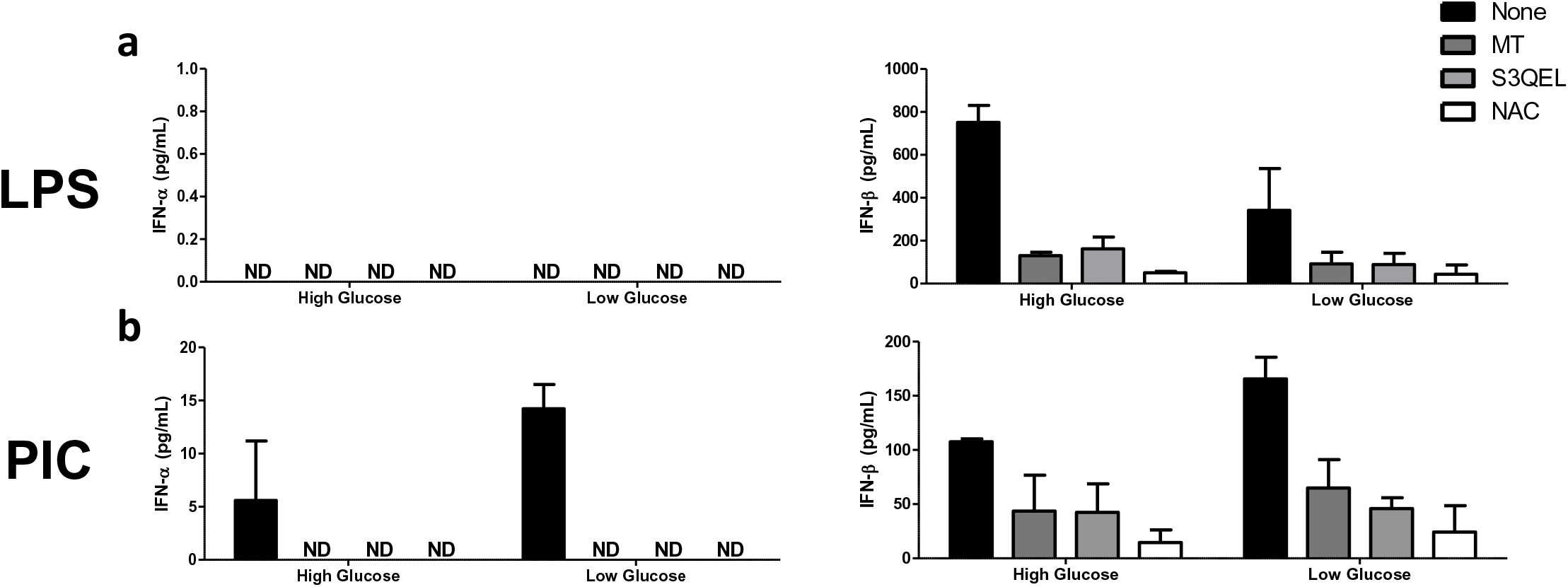
Targeting mitochondrial ROS leads to loss of type I IFN production. LPS-**(a)** or PIC-**(b)** stimulated BMDMs were co-treated with a panel of mtROS (MT, S3QEL, NAC) modulators to assess the importance of mitochondrial function to antiviral responses. CXCL10 and TNF-α cytokine secretion was measured after 18 hours in high glucose or low glucose media conditions. Data shown represents a mean of two individual animals.

